# Low Symbiodiniaceae diversity across Acropora hosts on the Central Pacific atoll Kiritimati

**DOI:** 10.64898/2026.10.08.757676

**Authors:** M. de Breuyn, A. Van Nynatten, J. K. Baum

## Abstract

The transformative impacts of global bleaching events are now a ubiquitous feature of coral reefs, often resulting in taxonomically simplified coral assemblages that can persist long-term due to ongoing ocean warming, heatwaves, and localized anthropogenic stressors. Understanding to what extent these processes shape symbiont biodiversity continues to be a major knowledge gap. Here, we examined *Acropora* host and symbiont communities on Kiritimati (Republic of Kiribati), seven and a half years after the 2015–2016 El Niño-associated marine heatwave, which caused a near total loss of the genus in the atoll’s shallow reefs. We quantified *Acropora* cover and growth-form composition at 37 sites spanning a gradient of local human disturbance and characterised Symbiodiniaceae communities using ITS2 DNA metabarcoding from 109 coral tissue samples at 13 of the sites. Corymbose and tabular *Acropora* growth forms remained evenly represented across the atoll and cover declined with increasing human disturbance. Symbiodiniaceae communities were dominated by *Cladocopium* with 93 % of colonies associated with a single ITS2 profile. Communities were homogenous, with low diversity and insignificant differences amongst growth forms, but slight differences across disturbance levels. DNA barcoding of the PaxC intron identified the majority of host colonies as most similar to *Acropora granulosa* (62 %) and *Acropora hyacinthus* (24 %). Although dominance by a single symbiont lineage is common among *Acropora* species across the Pacific, the low Symbiodiniaceae diversity observed at Kiritimati contrasts with the substantial within- and among-lineage diversity documented elsewhere. The depauperate Symbiodiniaceae communities observed in this study may thus represent a lasting legacy of the 2015–2016 disturbance, with the recovering *Acropora* assemblage representing an early stage of community reassembly drawing from a potentially depleted environmental symbiont pool.

## Introduction

Extreme climatic events (ECE’s) are increasingly recognized as a dominant force reshaping ecological communities across terrestrial, freshwater and marine ecosystems [1–5]. Marine heatwaves (MHWs), defined as periods of anomalously high sea surface temperatures, represent one of the most rapidly intensifying forms of ECE’s [6] and can drive rapid, large-scale mortality events that restructure community composition, often leaving assemblages that are less diverse and functionally simplified than those present prior to disturbance [4,7,8]. Because ecological recovery is typically uneven and species-specific, communities may follow alternative successional trajectories, with opportunistic or stress-tolerant taxa becoming disproportionately abundant while more sensitive species fail to re-establish [7,9–11]. These shifts can have lasting consequences for ecosystem function, particularly under repeated environmental stress events or when local anthropogenic stressors limit recovery to pre-disturbance levels [4,12,13]. Coral reefs are among the most climate-sensitive ecosystems and are expected to experience severe degradation under continued warming, given their sensitivity to even ∼1 °C increases in sea temperature [14]. Thermal stress disrupts the symbiosis between reef-building corals and their dinoflagellate algal partners (Symbiodiniaceae; [15,16]) leading to coral bleaching [17]. Extended bleaching periods frequently result in elevated mortality rates [18–20]. The increasing frequency and intensity of MHWs have intensified global mass bleaching events, with these acute thermal disturbances shaping bleaching severity, spatial extent, and subsequent reef recovery trajectories [6,18,21]. Recovery is further modulated by local anthropogenic stressors (e.g., pollution, fishing pressure, coastal development), which can impair resilience by altering nutrient regimes and destabilising symbioses [22,23], reducing calcification and physiological performance [24,25], and limiting reproductive output and larval supply [26,27].

Although the impacts of MHWs and mass coral mortality events on coral hosts are well documented [18,28,29], little is known about post-MHW dynamics of symbiont communities [30]. While bleaching can alter Symbiodiniaceae community composition, it remains less clear if these shifts are temporary or persistent. In some cases, post-bleaching recovery does not return corals to their original symbiont communities, instead leading to longer-term changes in composition and diversity [30–34]. These outcomes also appear to depend on ecological constraints, including the availability of compatible symbionts and host–symbiont specificity [35,36], while local anthropogenic disturbance can alter the composition and availability of Symbiodiniaceae across reef habitats [37].

Corals can associate with a diverse pool of Symbiodiniaceae but these associations are often host-specific, with different coral species showing affinities for particular Symbiodiniaceae lineages across broad geographic scales [38,39]. Coral–symbiont associations are established via vertical transmission, the inheritance from parent colonies, or horizontal symbiont acquisition from the environment during early life stages or after disturbance [40]. Symbiodiniaceae can also occur in free-living forms or environmental reservoirs [41], but these can differ substantially from those associated with coral tissue and mucus [42]. Symbiodiniaceae communities can shift with environmental conditions and thermal history [39,43], and are further structured by functional differences among taxa. For example, *Durusdinium* is generally associated with increased thermal tolerance, while *Cladocopium* is often linked to higher photosynthetic performance and growth, leading to shifts in dominance under thermal stress [43–45]. These trait differences contribute to differential persistence and reassembly of symbiont communities during post-bleaching recovery, where community composition may reflect both environmental filtering and host–symbiont specificity [34].

Among reef-building corals, the genus *Acropora* is particularly important for understanding symbiont dynamics under thermal stress due to its reliance on horizontal acquisition [40] and because it is often among the fastest recovering coral taxa following bleaching events [46,47]. This transmission mode means that post-bleaching recovery depends heavily on environmental symbiont pools, making *Acropora* a useful system for assessing how disturbance reshapes Symbiodiniaceae diversity and community composition [15,48]. *Acropora* are characterised by rapid growth and early sexual maturity [49,50], traits that increase susceptibility to thermal stress. However, these same traits can also support relatively rapid post-disturbance recovery, particularly where larval supply enables effective recruitment and recolonization [46,47,51,52]. The genus *Acropora* spans diverse morphologies (e.g., tabular, corymbose, digitate), contributing substantially to habitat complexity and reef accretion across reef systems [53,54], and occupies a wide range of ecological niches across depth and exposure gradients [47]. *Acropora* hosts diverse Symbiodiniaceae communities, commonly dominated by the genera such as *Cladocopium* and *Durusdinium*, with community composition often species specific, varying spatially and environmentally, and shifting in some species toward increased *Durusdinium* dominance post disturbance [15,44,55,56]. Despite this symbiont flexibility, recovery following bleaching is highly variable and appears constrained by local symbiont availability and environmental conditions [57,58].

During the 2015–2016 El Niño, *Acropora* experienced some of the most severe declines globally, with ∼33% loss of live tissue cover in the Caribbean [59], 41.5 % whole-colony mortality in Japan [60], 86% mortality in the Chagos Archipelago [61], and near-total loss of coral cover on Kiritimati, Republic of Kiribati [20]. Among these regions, Kiritimati experienced a MHW of globally unprecedented duration (∼ one year; [20]). Seven and a half years after this MHW, recovery on Kiritimati remains heterogeneous: with some species, and the symbionts they hosted pre-MHW, showing evidence of recovery while others remain largely absent and potentially locally extinct [30,62]. Recent work on Kiritimati highlights contrasting recovery trajectories, including *Platygyra ryukyuensis*, a stress-tolerant, horizontally transmitting coral that shows comparatively high post-disturbance persistence and recolonisation via environmental symbiont acquisition [34], and competitively dominant, vertically transmitting taxa such as *Pocillopora grandis* and *Montipora aequituberculata*, which recover primarily through survival of remnant colonies and local reproduction but exhibit more constrained flexibility in symbiont reassembly following bleaching [30]. *Acropora* differs in combining horizontal symbiont acquisition [40] with high ecological and competitive dominance [63,64], implying that recovery may be constrained by environmental availability of symbiont lineages. This combination makes the genus a key system for testing whether prolonged thermal stress leads to persistent losses or compositional restructuring of symbiont diversity in reef-building corals.

Our objectives were to assess *Acropora*–Symbiodiniaceae associations in the equatorial central Pacific since the 2015–2016 El Niño, establishing a new baseline against which the impacts of subsequent MHWs can be evaluated, and to assess whether *Acropora* cover and Symbiodiniaceae community composition vary with local human disturbance. We hypothesised that environmental filtering along the disturbance gradient would favour Symbiodiniaceae taxa tolerant of local conditions, producing both site-associated and shared assemblages, and that host-specific filtering would generate differences in symbiont composition among *Acropora* species [65,66]. We further predicted that *Acropora* cover would decline with increasing human disturbance, as chronic local pressures are associated with reduced coral cover and increased coral mortality by degrading habitat and intensifying environmental stress [20,67,68]. To address these questions, we characterised *Acropora* cover and Symbiodiniaceae communities across a gradient of local disturbance on Kiritimati, Republic of Kiritibati, the world’s largest coral atoll, where an unprecedented MHW (2015–2016) triggered a near total loss of the genus *Acropora*.

## Methods

### Study site and design

This research was conducted in the central equatorial Pacific Ocean on the remote atoll Kiritimati (Christmas Island; 01°52′ N, 157°24′ W), Republic of Kiribati. Kiritimati provides a unique natural setting for studying how human activities, including coastal development, pollution, and fishing, affect the resilience of coral reefs due to a clear gradient of human disturbance [69,70]. The atoll had an estimated human population of ∼7,400 in 2020 [71], with subsistence fishing representing a primary source of food security and livelihoods for much of the population [72]. Human activity is concentrated along the northwest coast, where approximately three-quarters of the island’s population resides [70,73], creating a clear gradient of local chronic human disturbance across the atoll (Fig 1). This gradient, quantified using population density and fishing pressure [37] and updated here with the most recent settlement data [73], is associated with substantial variation in coral cover and reef structural complexity across the atoll [69], despite largely consistent thermal environments [74]. Kiritimati’s coral reefs were severely impacted by the 2015–2016 El Niño MHW, which resulted in a ∼89% decline in hard coral cover at shallow forereef sites around the atoll. *Acropora* was among the most severely affected genera, with complete loss of all corymbose and tabular colonies reported at very low-disturbance sites [20]. We conducted this study via SCUBA in August 2023 at 37 long-term monitoring sites on Kiritimati’s shallow (10-12 m) forereef [20,75], with coral samples for genetic analyses collected from a subset of 13 of these sites (Fig 1). Sites were selected to encompass the full anthropogenic disturbance gradient, from reefs adjacent to the densely populated northwestern settlements to remote reefs in the sparsely inhabited southeastern region of the island, where human influence is minimal due to limited infrastructure and high transportation costs.

**Figure 1.**
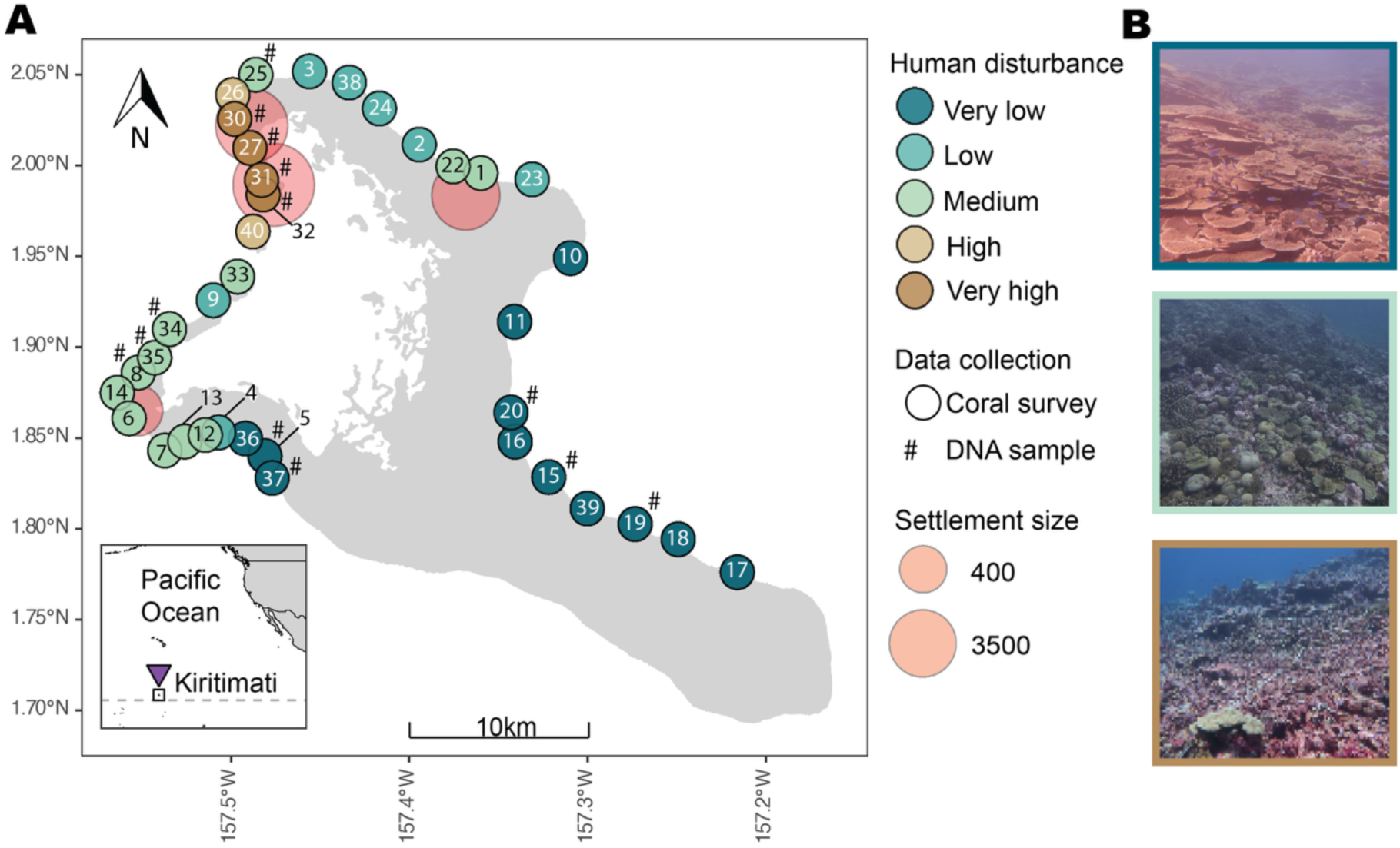
Kiritimati (Christmas Island) with forereef sites surveyed in 2023. (A) Distribution of the 37 surveyed sites (circles) and 13 sampling locations (#) along a gradient of human disturbance ranging from very low (blue) to very high (brown). Permanent human settlements are shown as red circles, with symbol size representing population size. (B) Visualisation of very low, medium, and very high disturbance sites.

### Coral surveys and sample collection

Surveys of benthic community composition were conducted at each site by placing a 1 m^2^ quadrat at fifteen randomly selected spots along two 25 m transects (separated by 10 m) positioned along the 10–12 m depth isocline. Each quadrat was photographed using a Canon EOS Rebel SL1 in an Ikelite housing, with white balance calibrated at depth before imaging. A total of 555 quadrat images were processed in CoralNet following the methods of Williams et al. [76]. Prior to upload, images were cropped to include only the quadrat area. Within CoralNet, 100 stratified random points were manually annotated per image using a custom benthic label set developed by Baum et al. [20] with organisms being identified to the lowest possible taxonomic level. *Acropora* was categorised by growth-form morphology, including tabular, corymbose, digitate, and *Acropora* spp. for recruits too small to assign to a specific growth form. Percent coral cover at each site was calculated to assess *Acropora* dominance by summing point counts for each growth form across all quadrats and expressing them relative to the total identifiable substrate points recorded at the site.

Coral samples for host DNA barcoding and symbiont DNA metabarcoding were collected at 13 of the 37 sites where previous coral tagging and sampling has been concentrated [30,34], restricted to very low, medium, and very high disturbance, along the same 50 m transects used for coral surveys. Sampling included tabular and corymbose *Acropora* colonies ranging from 15–55 cm in diameter (mean ± SD: 26.44 ± 7.73 cm) collected across a depth range of 7.8–12.8 m (11.03 ± 1.19 m). Colonies were spaced >5 m apart to reduce the likelihood of sampling the same genet [77] and collected at the upper central region of each colony to minimise potential bias arising from differences in microbial composition across intra-colony locations [78]. Fragments were broken off by hand or using a woodworking chisel, which was cleaned with gloves to remove residual tissue between sampling. Samples were placed in individual sterile bags and kept on ice until the end of the dive day. Coral samples were processed the same night by removing as much tissue as possible from the aragonite skeleton using an airbrush (Iwata-Medea Eclipse HP BCS) with compressed air and filtered seawater as the extraction medium. The resulting tissue slurry (∼0.3–0.6 μL) was collected and transferred into microcentrifuge tubes containing 400 μL of guanidinium isothiocyanate buffer. All samples were then stored at 4 °C until DNA extraction was performed.

### DNA extraction

To classify host species and assess algal symbiont diversity, a total of 109 *Acropora* colonies were processed for PaxC DNA barcoding and ITS2 DNA metabarcoding. DNA was extracted using a guanidinium-based method optimised for *Symbiodiniaceae* [79]. Briefly, 200 μL aliquots of coral tissue samples buffered in guanidinium, were incubated for 10 minutes at 72 °C and centrifuged to remove debris. DNA in supernatant was then precipitated overnight in 200 μL isopropanol at -20 °C. The sample was then centrifuged, and the DNA pellet was washed three times with 70% ethanol before re-suspending in 0.1M Tris buffer (pH 8). Extracted DNA was stored at −20°C until further processing.

### DNA barcoding of coral hosts

*Acropora* host species identities were attained by sequencing the PaxC intron of 82 samples. Amplifications were performed using GoTaq DNA polymerase (Promega) following the manufacturer’s protocol and *Acropora*-specific primers for the PaxC FP1 (5′-TCC AGA GCA TTA GAG ATG CTG G-3′) and RP1 (5′-GGC GAT TTG AGA ACC AAA CCT GTA-3′) [65] under the following thermocycler conditions: initial denaturation at 94°C for 4 min, followed by 35 cycles of 94°C for 30 s, 54°C for 30 s, and 72°C for 60 s, with a final extension at 72°C for 5 min. Successful amplification was verified by visualisation on a 1% agarose gel. PCR products were purified using ExoSapIT (Thermo Fisher Scientific) and Sanger sequenced at The Centre for Applied Genomics (TCAG), Toronto, Canada. Raw sequences were visualised, manually trimmed for quality, and assembled de novo in Mesquite [80]. Species assignments were made by querying PaxC sequences against a curated reference sequence database of Pacific *Acropora* species (Table S1) using BLASTn [81], with identity determined by the closest match to a reference sequence by bitscore. Representative sequences were aligned to the reference sequence dataset of Pacific *Acropora* PaxC sequences using MAFFT [82]. Alignment columns with more than 50 % gaps were removed. Phylogenetic relationships were inferred using maximum likelihood in IQ-TREE2 [83] with node support assessed via ultrafast bootstrap approximation. The resulting tree was visualized with ggtree [84].

### DNA metabarcoding of algal symbionts

Symbiodiniaceae ITS2 sequences were amplified and prepared for multiplex amplicon sequencing using a single-step library preparation approach with the SYM_VAR_5.8S2 and SYM_VAR_REV primers [85]. Each PCR reaction included 12.5 uL of Q5 HotStart PCR mastermix (New England Biolabs), 9.25 uL molecular grade water, 1 uL of each primer, and 1.25 uL of template DNA and was amplified using the following thermocycler conditions: initial denaturation at 98 °C for 2 min, followed by 35 cycles of 98 °C for 10 s, 56 °C for 30 s, and 72 °C for 30 s, with a final extension at 72 °C for 5 min. Sequencing was then carried out on the Illumina MiSeq platform using 2 × 250 bp paired-end chemistry. SymPortal (v0.3.24) [86] was used to group and merge ITS2 amplicon sequence variants (ASVs) into ITS2 type profiles by identifying consistent and recurring patterns in ASV occurrence and relative read abundance (RRA) across samples. These ITS2 type profiles provide a more ecologically meaningful and taxonomically robust representation of Symbiodiniaceae diversity than individual multicopy ITS2 ASVs generated by metabarcoding approaches [87]. Samples with fewer than 1000 sequencing reads were not included in the analysis.

### Statistical analyses

Image analyses were restricted to corymbose and tabular colonies. Digitate *Acropora* and *Acropora* spp. annotations were excluded from all analyses because they were not sampled for symbiont analysis due to low colony availability (digitate *Acropora*) and insufficient colony size (*Acropora* spp.). Monotonic trends in *Acropora* cover across the ordered disturbance gradient was assessed using Jonckheere–Terpstra tests with permutation-based p-values (10,000 permutations). Differences in percent substrate cover among disturbance categories and growth forms were assessed using Wilcoxon rank-sum tests. Visualisation was done with packages ggplot2 [88] and ggh4x [89].

Symbiont community composition was characterised using the ITS2 type profiles and UniFrac distances generated by Symportal [86]. Indicator species analysis was performed using the IndVal method in the indicspecies package [90] to identify taxa (ITS2 type profiles) associated with host growth form and disturbance. Weighted UniFrac distances of *Cladocopium* were calculated using Symportal [86] and the vegan package [91] to quantify phylogenetic dissimilarity among symbiont profiles following the UniFrac approach [92] and visualised using principal coordinates analysis (PCoA) with ggplot2 [88]. Differences in *Cladocopium* community composition across disturbance categories and host growth forms were tested using PERMANOVA (adonis2; vegan) [91] on weighted UniFrac distance matrices. Multivariate dispersion was assessed using betadisper and permutest (PERMDISP; vegan) [91] to evaluate differences in distances to group centroids among groups. Pairwise weighted UniFrac distances were used to compare within-host *Cladocopium* community variation across disturbance categories and host growth form using Wilcoxon rank-sum tests. P-values were adjusted for multiple comparisons using the Benjamini–Hochberg false discovery rate correction. All analyses were conducted in RStudio (v4.5.3) [93], with data manipulation performed using *dplyr [94], tidyverse [95]*, and *purrr* [96].

### Inventory of Symbiodiniaceae diversity in Pacific Acropora

To determine whether Symbiodiniaceae diversity of *Acropora* samples in this study was comparable to that reported from other Pacific regions, we compiled a database of published datasets available through the SymPortal database [86]. Studies were selected based on their focus on *Acropora* and restriction to Pacific reef systems. The profiles.relative.abund_and_meta and seqs.relative.meta_only outputs were downloaded for each study and used for subsequent comparison. ITS2 profiles were then aggregated into higher-level groups based on dominance (‘C1’, ‘C3’, ‘C40’, ‘C50’, ‘D’, ‘A’, and ‘others’), and percentages were calculated by dividing the total relative read abundance within each group by the total number of relative abundances across all groups and multiplying by 100. A summary table was generated to enable cross-study comparison of Symbiodiniaceae community composition.

## Results

### Acropora cover across disturbance

Total substrate cover of tabular *Acropora* was highest at very low disturbance sites (0.46 %) and declined sharply with increasing disturbance, reaching lows of 0.01 % and 0.00 % at high and very high disturbance sites, respectively. Corymbose *Acropora* had a similar pattern with highest mean substrate cover at very low disturbance sites (0.26 %), declining gradually to 0.01 % and <0.001% at high and very high disturbance sites, respectively (Fig 2A). This decline was supported by Jonckheere–Terpstra tests, which showed that both corymbose *Acropora* (JT = 115, p = 0.0001) and tabular *Acropora* (JT = 133, p = 0.0003) cover decreased significantly with increasing disturbance. The relative distribution of the two growth forms did not differ across disturbance levels (Wilcoxon rank-sum test, W = 829.5, p = 0.108).

**Figure 2.**
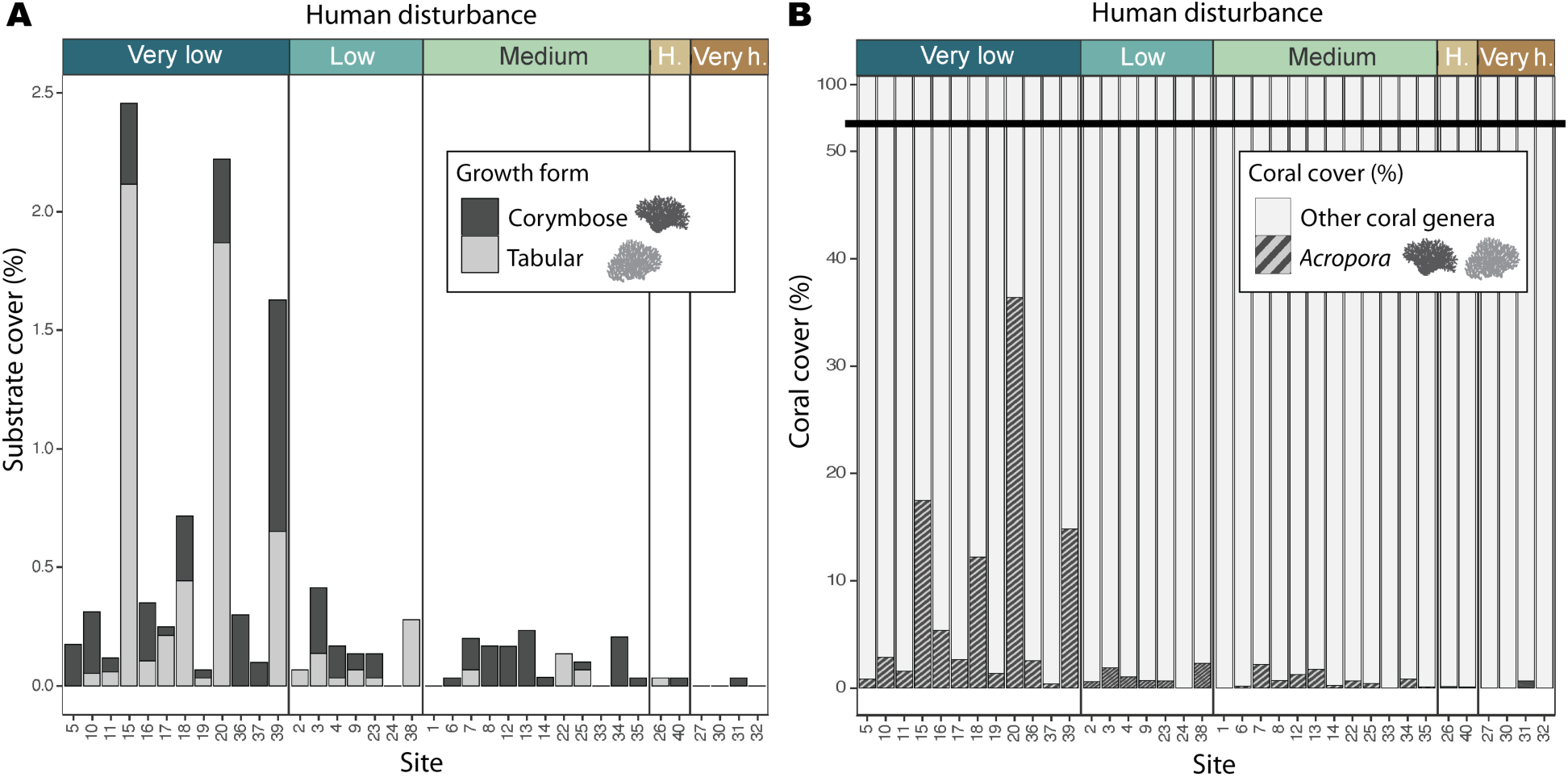
*Acropora* substrate and coral cover across a human disturbance gradient on Kiritimati in 2023. (A) Percentage substrate cover of corymbose *Acropora* (dark grey) and tabular *Acropora* (light grey). (B) Percentage coral cover of *Acropora* (corymbose and tabular *Acropora*; striped) and all other coral genera (white grey).

*Acropora* corymbose and tabular combined accounted for a small proportion of overall coral cover, with the highest cover recorded at very low disturbance sites with 8.2 %. This declined gradually to 0.6 % at high disturbance sites (99.4 % other coral genera), and 0.2 % at very high-disturbance sites (99.8 % other coral genera). The highest *Acropora* coral cover was recorded at site 20 (36.4 %), followed by site 15 (17.5 %), site 39 (14.9 %), and site 18 (12.2 %). These sites were all located close to one another within the very low human disturbance category (Fig 2B).

### Acropora Symbiodiniaceae diversity

Across all *Acropora* samples, symbiont communities were dominated by a small number of ITS2 profiles. Most colonies harboured a single dominant ITS2 profile, with 96.3 % of colonies assigned to one profile and only 3.7 % containing two or more profiles. In total, 109 colonies were assigned to nine profiles within *Cladocopium* (C), and 4 colonies assigned to two *Durusdinium* (D) profiles. The most prevalent profile was C1-C1c-C1b-C3-C42.2, detected in 50 colonies (38 corymbose, 12 tabular), followed by C1/C3-C1c-C1b, which occurred in 32 colonies (16 corymbose, 16 tabular). Other common profiles included C1-C1b-C1c-C42.2-C1bh (15 colonies: 14 corymbose, 1 tabular), C1-C1c-C1al (5 colonies: 3 corymbose, 2 tabular), and C1-C40-C1c (4 colonies: 2 corymbose, 2 tabular). The remaining four profiles accounted for two or fewer colonies. The thermotolerant *Durusdinium* [97] was rarely detected, with D1 and D4 each occurring in 2 colonies of tabular *Acropora* and *Durusdinium* being the dominant symbiont in only one of these (Fig 3).

**Figure 3.**
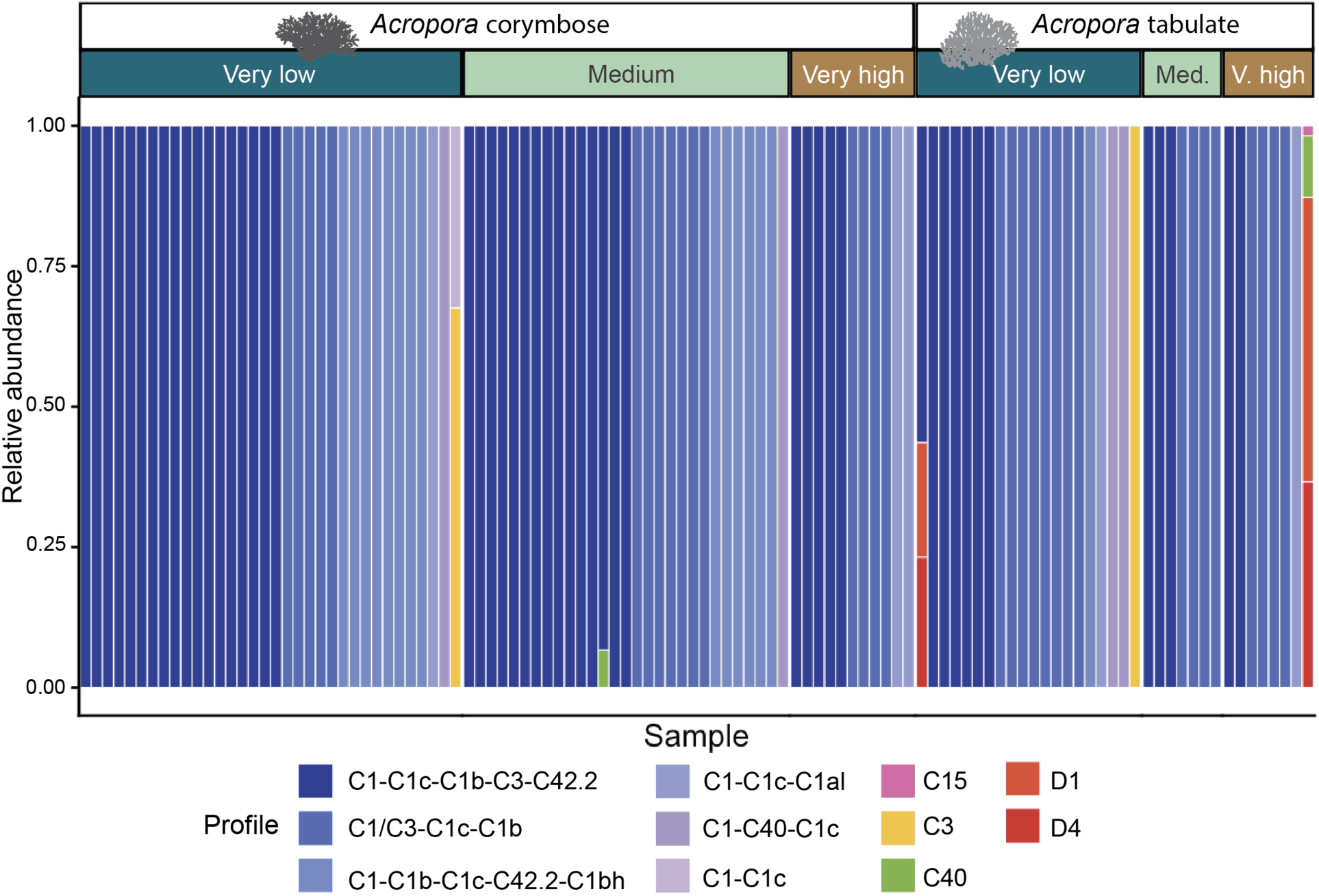
Relative abundance of ITS2 profiles in 109 Acropora corals across growth forms (corymbose, tabular) and levels of exposure to chronic local human disturbance. Each bar represents an individual coral sample, with colours indicating the relative abundance of *Cladocopium* (C; shades of blue – purple, pink, yellow, green) and *Durusdinium* (D; orange, red) profiles.

Symbiodiniaceae profile composition and relative abundance were broadly similar among growth forms and disturbance gradients, with considerable overlap observed across most samples. PERMANOVA analyses detected significant effects of disturbance level (*R*² = 0.0267, *F* = 2.93, *p* = 0.019) and host growth form (*R*² = 0.0303, *F* = 3.34, *p* = 0.010) on *Cladocopium* community composition, although these factors explained only a very small proportion of the total variation (2.67 % and 3.03 %, respectively). Consistent with these low effect sizes, PCoA ordination showed substantial overlap among disturbance and no clear separation by growth form (Fig 4A), despite slightly greater dispersion in ordination space of tabular *Acropora* compared to corymbose.

**Figure 4.**
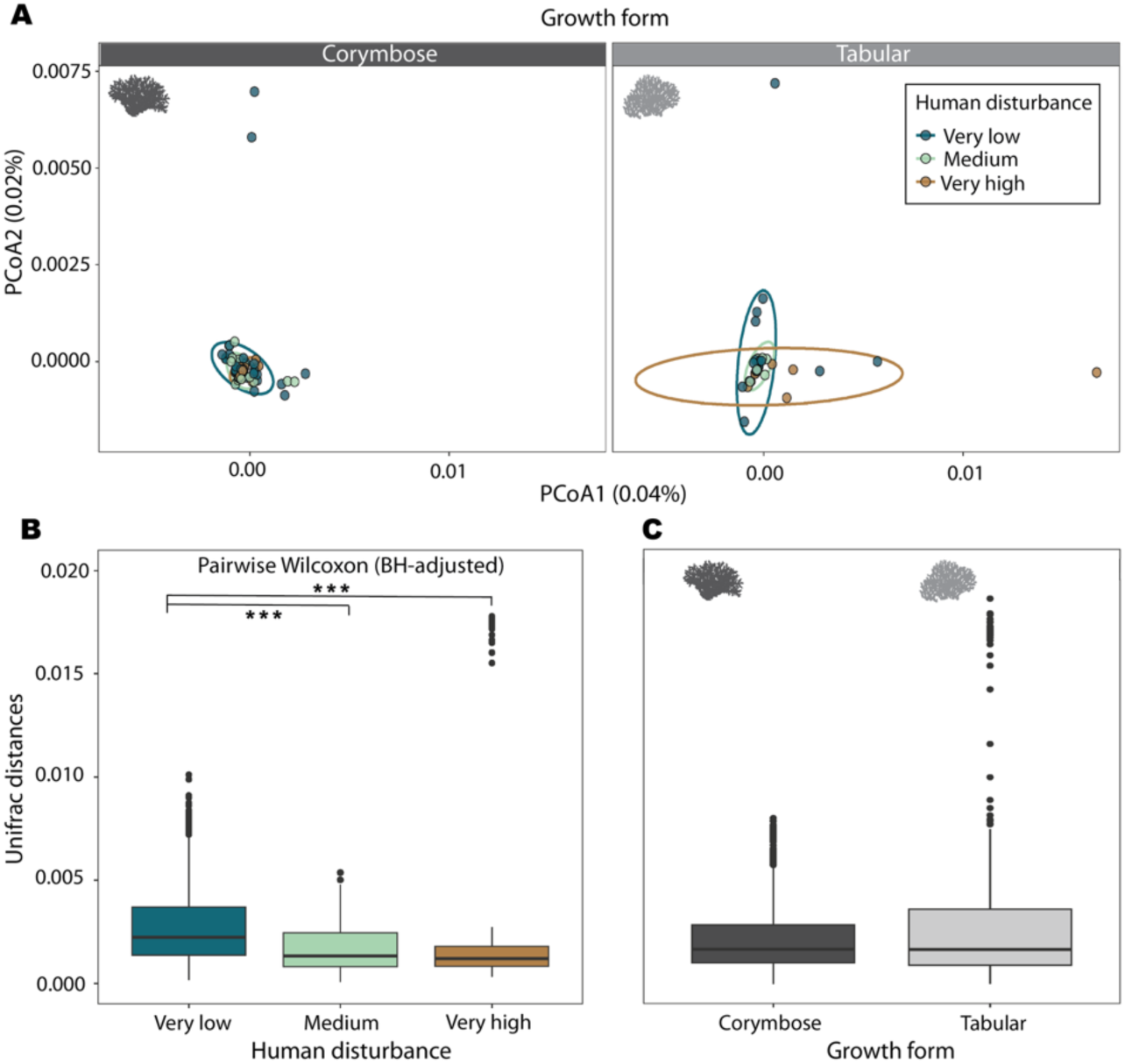
Cladocopium community composition across *Acropora* hosts and disturbance gradient. (A) Principal Coordinates Analysis (PCoA) based on Unifrac distances of Cladocopium ITS2 assemblages associated with *Acropora* corymbose and tabular colonies across three disturbance levels (very low, medium, and very high). Comparison of Cladocopium community composition based on UniFrac distances amongst *Acropora* colonies (B) collected from sites exposed to different disturbance levels, (C) within the same growth form (corymbose (left) and tabular (right)). *** indicate p < 0.001.

Pairwise Wilcoxon rank-sum tests (with BH correction) revealed significant differences among disturbance levels. Dispersion, measured as the distance of samples from their group centroid in UniFrac space, differed significantly between very low and medium disturbance (*p* < 2 × 10⁻¹⁶) and between very low and very high disturbance levels (*p* < 2 × 10⁻¹⁶), but not between medium and very high disturbance (*p* = 0.20; Fig 4B).

When grouped by growth form, multivariate dispersion was greater in tabular *Acropora* (mean distance to centroid = 0.00191) than in corymbose *Acropora* (0.00143). Within-host pairwise distances (Wilcoxon rank-sum test, *p* = 0.989; Fig. 4C) did not differ significantly between corymbose and tabular *Acropora*. Indicator species analysis revealed limited host specificity among ITS2 profiles. The profile C1–C1b–C1c–C42.2–C1bh was significantly associated with corymbose *Acropora* (IndVal = 0.421, *p* = 0.021), whereas C1–C3–C1c–C1b was significantly associated with tabular *Acropora* (IndVal = 0.567, *p* = 0.0045). Specificity across the human disturbance gradient was similarly weak, with only one profile, C1–C1c–C1al, identified as a significant indicator of the very high disturbance category (IndVal = 0.372, *p* = 0.0078).

### Species identification of Acropora growth forms

Classification of *Acropora* PaxC barcode sequences from a subset of colonies (n = 82) revealed, that the majority of tagged corals were most similar to *A. granulosa* (Milne Edwards, 1860; n = 51) and *A. hyacinthus* (Dana, 1846; n = 21), largely aligned with colony growth forms of corymbose and tabular (Table S2). Maximum likelihood phylogenetic reconstruction of representative samples from corymbose and tabular colonies further supports these relationships (Fig 5A & B). Seventeen colonies did not conform to these dominant growth form to referencereference species associations. One tabular colony was most similar to *A. granulosa* and 5 corymbose were most similar to *A. hyacinthus*. An additional 10 samples had the highest sequence similarity to *Acropora digitifera* (7 corymbose), *Acropora muricata* (1 tabular), and *Acropora selago* (2 tabular; Table S2). Both *A. granulosa* and *A. hyacinthus* are previously undocumented in the Line Islands based on species ranges in online resources Corals of the World and World Register of Marine Species (WoRMS) [98,99], and no other *Acropora* coral barcode sequences are available for Kiritimati.

**Figure 5.**
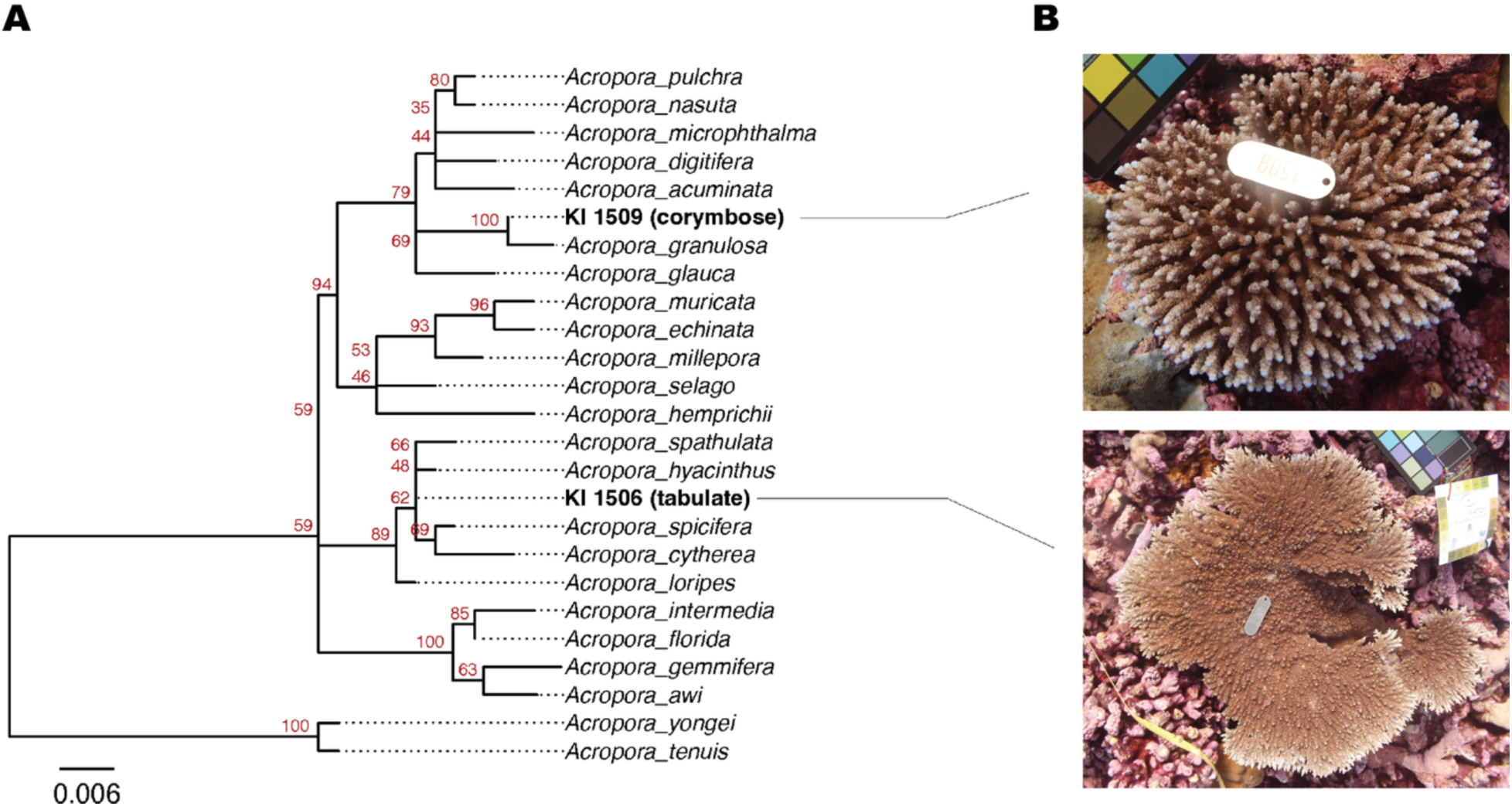
PaxC phylogeny and representative *Acropora* taxa. (A) Phylogenetic reconstruction of representative PaxC sequences from predominant *Acropora* species on Kiritimati. Bootstrap support for each node indicated in red. (B) Images of representative taxa.

### Inventory of Acropora symbiont diversity across the Pacific

Comparison of Symbiodiniaceae profiles within *Acropora* across eight published SymPortal datasets [87] revealed differences in the dominant ITS2 profiles among Pacific island locations and *Acropora* species (Table 1). *A. hyacinthus* showed the greatest variation among datasets, with C3 dominant in the Great Barrier Reef (GBR) in 2020 (56 %) [100] and French Polynesia in 2013 (94 %) [101], while *Symbiodinium* was dominant in another French Polynesian dataset in 2019 (60 %) [102]. None of which match the C1 that was dominant in *A. hyacinthus* from the current 2023 Kiritimati dataset (92 %). Substantial variation was seen in the symbiont profiles dominant in other published *Acropora* datasets. C40 was the dominant profile across all three *A. digitifera* datasets from Palau, sampled in the years 2018, 2019, and 2022, accounting for > 99 % of relative ITS2 sequence abundance [103–105]. In New Caledonia, C1 dominated *Acropora muricata* datasets sampled in 2016 and 2021 (87 % and 51 %, respectively) [106,107], whereas *A. pulchra* was dominated by A (*Symbiodinium*) in 2016 (55 %) [106]. Overall, C1, C3, C40, and *Symbiodinium* were each the dominant profile in at least one of the eight datasets [100–107], whereas D accounted for <1 % of relative ITS2 sequence abundance in three datasets [104–106] and 12 % in one dataset [102]. To the best of our knowledge, no data for symbiont communities of *A. granulosa* was available on Symportal.

**Table 1.** Descriptive inventory of *Acropora* symbiont availability across the Pacific by major Symbiodiniaceae clade (C1, C3, C40, C50, D, A, and others), based on total ITS2 profile relative read abundance. Dominant symbiont groupings (%) are highlighted in yellow, and studies are ordered by *Acropora* species from most recent year sampled to earliest. The current study is shown in bold.

| Study | *Acropora* species | Location | Depth | Year sampled | Total ITS2 profile sample abundance |  |  |  |  |  |  | Total ITS2 profile relative read abundance (%) |  |  |  |  |  |  |
| --- | --- | --- | --- | --- | --- | --- | --- | --- | --- | --- | --- | --- | --- | --- | --- | --- | --- | --- |
|  |  |  |  |  | C1 | C3 | C40 | C50 | D | A | others | C1 | C3 | C40 | C50 | D | A | others |
| de Breuyn et al. | *A. granulosa* | Kiritimati, Kiribati | 8-13 | 2023 | 51 | 1 | 1 | 0 | 0 | 0 | 0 | 98.55 | 1.32 | 0.13 | 0 | 0 | 0 | 0 |
| de Breuyn et al. | *A. hyacinthus* | Kiritimati, Kiribati | 8-13 | 2023 | 19 | 0 | 1 | 0 | 2 | 0 | 1 | 92.82 | 0 | 0.55 | 0 | 6.54 | 0 | 0.09 |
| Buzzoni et al. 2026 | *A. digitifera* | Palau | 3-8 | 2022 | 0 | 0 | 295 | 2 | 0 | 0 | 13 | 0 | 0 | 99.39 | 0.32 | 0 | 0 | 0.29 |
| Humane s et al. 2022 | *A. digitifera* | Palau | 0-4 | 2019 | 0 | 0 | 97 | 0 | 2 | 0 | 0 | 0 | 0 | 99.81 | 0 | 0.19 | 0 | 0 |
| Lachs et al. 2023 | *A. digitifera* | Palau | 1-3 | 2018 | 0 | 0 | 51 | 0 | 1 | 0 | 0 | 0 | 0 | 99.36 | 0 | 0.64 | 0 | 0 |
| Grima et al. 2022 | *A. hyacinthus* | GBR, Australia | 2-5 | 2020 | 1 | 9 | 1 | 1 | 0 | 0 | 5 | 1.21 | 56.81 | 11.07 | 5.79 | 0 | 0 | 25.12 |
| Leinbach et al. 2023 | *A. hyacinthus* | Moorea, French Polynesia | 3-14 | 2019 | 39 | 67 | 0 | 0 | 38 | 102 | 1 | 8.58 | 17.69 | 0 | 0 | 12.96 | 60.77 | 0 |
| Kriefall et al. 2022 | *A. hyacinthus* | Moorea, French Polynesia | 3-6 | 2013 | 0 | 90 | 0 | 0 | 0 | 9 | 0 | 0 | 94.03 | 0 | 0 | 0 | 5.97 | 0 |
| Alessi et al. 2024 | *A. muricata* | New Caledonia, France | NA | 2021 | 30 | 4 | 0 | 26 | 0 | 0 | 3 | 51.01 | 4.74 | 0 | 40.95 | 0 | 0 | 3.29 |
| Alessi et al. 2024 | *A. tenuis* | New Caledonia, France | NA | 2021 | 2 | 1 | 0 | 0 | 0 | 0 | 0 | 61.83 | 38.17 | 0 | 0 | 0 | 0 | 0 |
| Camp et al. 2020 | *A. muricata* | New Caledonia, France | 1-3 | 2016 | 7 | 0 | 0 | 1 | 1 | 0 | 0 | 87.23 | 0 | 0 | 12.34 | 0.43 | 0 | 0 |
| Camp et al. 2020 | *A. pulchra* | New Caledonia, France | 1-3 | 2016 | 1 | 7 | 0 | 5 | 2 | 5 | 1 | 0.56 | 36.52 | 0 | 5.99 | 0.64 | 55.31 | 0.98 |

## Discussion

We found a highly homogeneous community composition of algal symbionts hosted by *Acropora* across Kiritimati, despite substantial variation in host species abundance, growth forms, and ecological conditions across the atoll. This striking lack of spatial differentiation may represent a lasting legacy of the severe 2015–2016 MHW, with subsequent *Acropora* recovery likely occurring from a substantially depleted environmental symbiont pool. These findings highlight the need for reef managers to look beyond visible coral loss and consider biodiversity at the coral holobiont level. We suggest that *Symbiodiniaceae* diversity should be incorporated into monitoring and intervention strategies, because recovery in coral cover alone may mask less visible losses of ecologically important *Symbiodiniaceae* diversity, with potential consequences for the resilience of recovering coral populations.

### Cladocopium dominance across Acropora hosts and disturbance

Symbiodiniaceae communities seven and a half years after the 2015–2016 MHW were consistently dominated by *Cladocopium* across *Acropora* growth forms and disturbance regimes, while *Durusdinium* was almost absent. This pattern aligns with observations from across Pacific reef systems, where adult *Acropora* typically host *Cladocopium*-dominated communities, particularly C1, C3 and C40 lineages, with *Durusdinium* generally occurring at low abundance or absent [101,103–105,107]. In contrast, other coral genera from Kiritimati have shown pronounced post-bleaching shifts toward the thermal resistant *Durusdinium* post MHW [30,32,34]. Claar et al. [32] reported transitions from *Cladocopium*-to *Durusdinium*-dominated communities in *P. ryukyuensis* and *Favites pentagona* between 2015 and 2017, while Buzzoni et al. [34] demonstrated that this state persisted for at least eight years in *P. ryukyuensis* with 92 % of colonies remaining dominated by *Durusdinium* and limited recovery of pre-MHW *Cladocopium* lineages. Similarly, Van Nynatten et al. [30] documented a widespread reorganisation of Symbiodiniaceae assemblages in *P. grandis* and *M. aequituberculata*, with sustained *Durusdinium* dominance for at least seven years post-disturbance.

The difference in symbiont dominance across genera is likely related to symbiont transmission mode. *Acropora* are predominantly horizontal transmitters [40], acquiring Symbiodiniaceae from the surrounding environment during early ontogeny. However, uptake is not necessarily proportional to environmental availability and can be influenced by host selectivity during early symbiosis [55,108]. Given the near-complete loss of *Acropora* during the 2015-2016 MHW, present-day colonies likely represent post-disturbance recruits that established symbioses from the altered environmental symbiont pool. Despite the increased prevalence of *Durusdinium* in other Kiritimati corals following the MHW [30,32,34], *Acropora* were predominantly associated with C1 lineages. The contrast between symbiont communities detected in *Acropora* and other coral genera from Kiritimati suggests that these recruits did not simply acquire the most prevalent environmental symbionts, but that host-mediated filtering may favour *Cladocopium* within *Acropora* [109,110]. The dominance of C1 in our *Acropora* assemblages may also be consistent with a favourable host–symbiont association, as *Acropora tenuis* juveniles hosting C1 have been shown to incur lower metabolic costs and exhibit greater tolerance to combined heat and light stress than those hosting type D [111].

### Low Symbiodiniaceae diversity across Acropora hosts

Symbiodiniaceae communities showed strong overlap across *Acropora* hosts, with low diversity both within individual colonies and across the symbiont assemblage. Nearly all colonies (96 %) were assigned to a single ITS2 profile, and the assemblage was dominated by mostly three recurring profiles, indicating a highly homogenised symbiont community across *Acropora* hosts. This contrasts with previous studies showing greater differentiation in Symbiodiniaceae communities among *Acropora* species. Camp et al. [106] found different dominant symbionts in *A. muricata* and *A. pulchra* in New Caledonia, while Quigley et al. [55] reported significant differences in symbiont composition among *A. hyacinthus*, *A. millepora*, and *A. tenuis* on the GBR, Australia. At Kiritimati, however, the same C1-associated profiles predominated across *Acropora* species, suggesting weaker host-specific structuring of Symbiodiniaceae communities at this location. This low diversity further contrasts with other Pacific *Acropora* populations, where strong dominance by a single symbiont lineage can coexist with greater diversity both within lineages and among lineages [100,102,103,107]. In French Polynesia, for example, *A. hyacinthus* was dominated by A-lineage symbionts (60 % of relative ITS2 profile abundance), but also hosted C1 (8 %), C3 (17 %), and D-lineages (∼13 %) [102]. In Palau, *A. digitifera* was dominated by C40-lineage (∼99 %), yet, considerable fine-scale diversity persisted within this dominant lineage, with a dozen C40-associated ITS2 sequences reported [103]. The comparatively low diversity of *Acropora* symbiont assemblages on Kiritimati may therefore reflect post-disturbance recolonisation from a simplified regional pool of available *Symbiodiniaceae*, likely reflecting the severity of the 2015–2016 MHW and associated widespread coral loss [20]. This event likely removed much of the pre-existing host and Symbiodiniaceae heterogeneity on Kiritimati. Notably, C31 is thought to have gone locally extinct following the MHW, as the loss of its coral host, *M. aequituberculata*, may have eliminated the habitat required for its persistence [30], and multiple *Cladocopium* lineages prevalent in *Platygyra* and *Porites* colonies before the MHW are now absent or in very low frequency across hosts [34,112]. Following severe MHWs, coral–Symbiodiniaceae associations can undergo substantial restructuring, with changes persisting for years after the initial disturbance and, in some cases, resulting in homogenisation of symbiont communities across environmental gradients [30]. Given the greater symbiont diversity and community differentiation reported from *Acropora* populations elsewhere in the Pacific [100–107], the strong homogenisation observed on Kiritimati may represent a lasting legacy of the 2015–2016 disturbance, with the subsequent recovery of *Acropora* occurring from a substantially depleted symbiont pool.

### Acropora species identity predominantly matches growth form

Overall, molecular-based species classifications were broadly consistent with morphology-based identifications, although several colonies did not conform to the dominant growth form–species associations. The majority of sampled colonies were most similar to *A. granulosa* (62 %) or *A. hyacinthus* (24 %) based on available reference sequences for *Acropora* species in the Pacific available on GenBank. Although *Acropora* species have previously been documented from Kiribati based on morphological characteristics [20,113,114], to the best of our knowledge, this study provides the first DNA-based species identification for the genus *Acropora* from Kiribati. Based on species ranges in Corals of the World and World Register of Marine Species (WoRMS) [98,99], two commonly used online resources for coral species identification and geographic distribution, our data provide previously undocumented records of both *A. granulosa* and *A. hyacinthus* from the Line Islands. *A. hyacinthus* was previously recorded on Tarawa (Gilbert islands) on WoRMS [115] and marked as strongly predicted for occurrence on Kiritimati on Corals of the World [116], whereas *A. granulosa* had no listed records or predictions [117,118]. Given the frequent mismatch between morphological and molecular species boundaries in *Acropora* [65,119], our barcode-based analyses provide a more accurate understanding of these species’ distribution across the Pacific. Understanding the current distribution of coral species is critical given how sensitive these organisms are to MHWs, which are increasing in frequency and intensity [6]. Accurate baseline distribution data is urgently needed to detect when and where populations of specific species are declining, contracting, or persisting under repeated heat stress.

### Limited *Acropora* coral cover recovery post mass mortality event

Despite the strong homogeneity of *Symbiodiniaceae* communities across *Acropora* hosts, the genus has persisted across the atoll since the near total loss after the 2015-2016 MHW, although recovery has remained limited. In comparison to coral cover estimates in 2015 (pre-MHW) and 2016 (post-MHW) in Baum et al. [20], recovery on Kiritimati was greatest at medium disturbance sites, where *Acropora* cover reached 0.1 % of total substrate cover in 2023, representing 13.1% (0.1/0.8 × 100) of the pre-MHW cover of 0.8 %. This was followed by very low disturbance sites, where cover increased from 0 % in 2016 to 0.7 % in 2023, but recovered to only 7.0 % (0.7/10.3 × 100) of its pre-MHW cover of 10.3 %. At very high disturbance sites, *Acropora* cover was already extremely low before the MHW (0.01 %) and remained almost absent in 2023 (<0.01 %). This slow recovery is consistent with studies showing that coral reefs can take more than a decade to recover from severe disturbance [46,120–123]. The near-total loss of *Acropora* post MHW [20] may have constrained recovery by reducing the local population available to replenish the reef. This may be particularly important on remote reefs, where limited connectivity can restrict larval supply following widespread mortality [46]. For example, despite a 95 % decline in *Acropora* cover following the 2016 and 2017 mass bleaching events at Lizard Island, a well-connected reef on the GBR, Australia [124], rapid recovery occurred where recruitment was high, with newly recruited colonies driving substantial increases in *Acropora* cover within just two years [52]. In contrast, following a 98 % decline in coral cover after the 2015–2016 MHW, the isolated Jarvis Island (365 km southwest of Kiritimati) [125] showed little coral recovery and shifted toward an algal-dominated state over the subsequent two years [126].

*Acropora* recovery also differed between growth forms. Corymbose *Acropora* generally showed greater recovery than the tabular growth form, particularly at very low- and medium-disturbance sites, where it regained a larger proportion of its pre-MHW cover despite having lower initial abundance. This suggests that recovery following the MHW may have been more constrained for tabular colonies than for corymbose colonies. This uneven recovery is consistent with previous studies showing that coral taxa and growth forms can recover at different rates following major disturbances, potentially leading to shifts in community composition during recovery [46]. Thus, the 2023 *Acropora* assemblage on Kiritimati may represent an early stage of community reassembly rather than a return to its pre-MHW composition.

### Acropora cover declined with disturbance, but growth forms remained consistent

*Acropora* cover declined strongly along the disturbance gradient, with the highest cover recorded at pristine reefs and *Acropora* becoming rare or absent at sites located close to human settlements. This pattern is consistent with previous findings on Kiritimati showing declines in coral cover with increasing anthropogenic disturbance [20] and reflects the well-documented sensitivity of *Acropora* to local stressors associated with coastal development, including sedimentation and nutrient enrichment [18,127].

Despite the pronounced decline in coral cover, the relative contribution of corymbose and tabular growth forms remained consistent across disturbance levels. This is unexpected because coral growth forms differ in their susceptibility to environmental stressors, with previous studies demonstrating morphology dependent responses of *Acropora* to thermal and physical disturbance [128,129]. Moreover, the two species are associated with contrasting hydrodynamic environments: *A. hyacinthus* typically dominate shallow, wave-exposed reef-crest habitat [63,130], whereas *A. granulosa* is more found on reef slopes protected with lower wave exposure [131]. Both habitat types occur across Kiritimati, including exposed reef crests in the east and more sheltered reef slopes along the northwestern atoll. The consistent growth-form distribution we observed may therefore suggest that human activity exerts comparable cumulative pressures on both morphologies, potentially contributing to morphological homogenisation despite their contrasting ecological associations. A similar pattern was reported by Ford et al. [68], who examined 62 outer reefs across 17 Central and Western Pacific Island nations and territories and found that higher local human impacts were associated with lower live coral cover, while the composition of coral communities showed comparatively limited changes in morphology.

### Limitations and future directions

As the first assessment of Symbiodiniaceae communities associated with *Acropora* on a central equatorial Pacific reef, this study provides an important baseline for future research. Interpreting our findings within a broader Pacific context is complicated by Kiritimati’s geographic isolation, with the nearest neighbouring island, Tabuaeran, ∼ 285 km away, and the spatial and temporal heterogeneity of connectivity across the Line Islands [132]. Larval dispersal among the Northern Line islands can vary substantially with island pair and dispersal period [133], with such variability in connectivity contributing to genetic differentiation among geographically isolated coral populations across the Pacific [134]. Consequently, patterns observed at Kiritimati may reflect local dispersal dynamics and environmental conditions that differ from those of other Pacific reef systems. In the absence of historical data for *Acropora* symbiont communities on Kiritimati, we cannot determine whether the associations observed here are stable through time or reflect more recent environmental conditions. Continued monitoring will therefore be important for assessing potential temporal variation in symbiont communities and confirming our result of low Symbiodiniaceae diversity in *Acropora* hosts across the atoll. Planned sampling through the 2026–2027 super El Niño [135] will provide a unique opportunity to examine how *Symbiodiniaceae* communities in *Acropora* respond to a major MHW and whether the current low symbiont diversity persists or shifts in response to the predicted heat stress event.

## Conclusion

Across Kiritimati, Republic of Kiribati, *Acropora* hosted homogenous algal symbiont communities despite differences in host composition, growth form, and local ecological conditions, seven and a half years after near total loss of the genus around the atoll. Our results suggest that the effects of severe heat stress events can persist well beyond the initial loss of corals, with consequences for the composition of recovering symbiont communities and the wider coral holobiont. Recognising biodiversity losses that are less visible to the human eye will be critical for understanding the long-term trajectory of coral reef ecosystems under climate change. It is urgent that effective interventions address the coral holobiont, protecting not only coral hosts but also the diversity of their symbionts and the biological partnerships that underpin reef functioning.

## Author contributions

Conceptualisation: M.D.B., A.V.N., J.K.B. Permits, Funding: J.K.B. Data processing: M.D.B., A.V.N., Data analysis: M.D.B., A.V.N. M.D.B. wrote the original draft of the manuscript and all authors reviewed and provided edits to the manuscript. All authors approved the final version of the manuscript for publication.

## Acknowledgements

We extend our gratitude to the Government of the Republic of Kiribati and the people of Kiritimati for their continued support and hospitality, which made this research possible. We acknowledge with respect that the University of Victoria is located on the traditional territories of the Lekwungen-speaking peoples, including the Songhees, Esquimalt, and WSÁNEĆ Nations, whose enduring relationships with these lands and waters continue to this day. We thank the 2023 Baum lab field team for their dedication and assistance in data collection. Research was conducted under permits issued by the Ministry of Fisheries, Republic of Kiribati (Permit No. 032023), and OB 3/83.

## Supporting information

**S1 Table. Curated reference sequence database of Pacific *Acropora* species.**

**S2 Table. PaxC-based species assignments of *Acropora* colonies from Kiritimati.**

